# Comparative Genome Study of Multidrug-resistant *Serratia marcescens* strain IU-BTGE-M-3 Isolated from a Diarrheal Patient Reveals Antibiotic Resistance Profile

**DOI:** 10.1101/2024.10.18.619060

**Authors:** Md. Shohorab Hossain, Md. Abdullah Al Mamun, Dalal Sulaiman Alshaya, KOTB A Attia, Changhee Han, AKM Ariful Haque, Md Mafizur Rahman

## Abstract

*Serratia marcescens* is a notable pathogen known for its intrinsic and acquired antimicrobial resistance, posing challenges in healthcare. The study investigates the multidrug-resistant (MDR) *Serratia marcescens* strain BTGE-M-3, focusing on its genomic features and antibiotic resistance. The strain was identified using both morphological and molecular approaches, and its genome, which measured 4.97 Mbp and exhibited a GC content of 59.7%, was sequenced. The genome has 4,827 genes, including essential antibiotic resistance genes, such as *tet* (41) for tetracyclines, *aac* (6′)-*Ic* for aminoglycosides, *OqxB* for fluoroquinolones, and *bla*SST-1 beta-lactam, alongside various virulence determinants. Comparative genomics revealed high similarity (98.06% ANI) to *S. marcescens* strain KS10, supporting its classification within the same species. Additionally, the genome analysis showed that *S. marcescens* strain IU-BTGE-M-3 comprises of 4390 gene clusters, 4564 genes, and 244 unique singleton genes. To summarize, the results showed that *S. marcescens* strain IU-BTGE-M-3 exhibited a high level of genomic diversity as well as diverse metabolic, cellular, and biological functions, and it is hypothesized that frequent strain exchanges resulted in the horizontal transfer of drug resistance genes. This study underscores the importance of genomic surveillance in understanding and combating antibiotic resistance in therapeutic settings.

## Introduction

*Serratia marcescens* is commonly found in human [1], animals [2], insects [3], water (damp bathroom)[4], soil [5], plants [6–8], and nosocomial (hospital) environments [9], emphasizing its versatility and potential for causing infections in diverse settings [4]. *S. marcescens* has drawn global attention as an emerging pathogen, with its pathogenicity in humans being identified in 1913. Since then, studies have discovered an increase in the frequency of conditions brought on by this organism [10–12].

*S. marcescens* exhibits robust intrinsic antimicrobial resistance and possesses a remarkable capacity for acquiring, transferring, and modifying antimicrobial resistance genes [13]. Traditionally, third-generation cephalosporins or fluoroquinolones were used to treat *S. marcescens* infections; despite this, the number of multidrug-resistant strains has increased, complicating therapeutic approaches[4,14]. Regardless of its relatively low virulence, *S. marcescens* produces nosocomial infections in very immunocompromised or severely sick patients, specifically in environments such as intensive care units (ICUs), especially neonatal intensive care units (NICUs)[13,15]. Additionally, *S. marcescens* is frequently implicated in hospital-acquired infections[16], with multidrug-resistant strains being common and linked to outbreaks[12]. This bacterium inherently exhibits resistance to various antibiotic families, which is further compounded by acquired and adaptive resistance mechanisms.

The overuse of antibiotics, often for viral infections like the common cold in South Asia, leads to expand of antimicrobial resistance (AMR) [17]. The over-prescription of antibiotics, often unnecessary, is a result of non-qualified healthcare providers and unregulated healthcare industries. The root cause of antibiotic resistance in most of Asian countries, including Bangladesh [17], lack of access to safe drinking water, poor sanitation, and/or inadequate sanitation, and the widespread accessibility of antibiotics sold over the counter, facilities contribute to the spread of antibiotic-resistant “superbugs [18].” The COVID-19 pandemic has further complicated matters, with reports of indiscriminate antibiotic use potentially fueling AMR [19]. Scientists are concerned about the rising rate of AMR in Bangladeshi populations [20]. In Bangladesh, hospitals have emerged as hotspots for AMR, with concerns about the increasing prevalence of resistant strains [20,21]. Third-generation cephalosporins are prescribed more frequently than other antibiotics in Bangladeshi hospitals (60%), with macrolides being administered more frequently (40%) [22]. By applying the disk diffusion method and the criteria of the Clinical and Laboratory Standards Institute (CLSI), the incidence of antibiotic resistance is very high from hospital-acquired infections in Bangladesh [23]. Despite the widespread use of phenotypic methods like disk diffusion, which can detect resistance, there is limited data available on the genetic foundation regarding antibiotic resistance in *S. marcescens* and other bacterial pathogens in this region. This gap underscores the need for comprehensive genomic studies to identify and characterize antibiotic resistance genes, which would provide a deeper understanding of resistance mechanisms and support more effective infection control strategies.

This study focused on *S. marcescens* and their antibiotic resistance profiling. Whole genome sequencing to obtain complete genetic information on *S. marcescens*. The isolated *S. marcescens* were identified from fecal samples of a diarrhea patient. The comparative study can provide insights into the underlying genetic mechanisms that contribute to resistance and as well as contributing to our greater understanding of antibiotic resistance in therapeutic settings and informing effective treatment options.

## Materials and Methods

### Sample collection

The stool sample was taken from a 10-month-old girl who was admitted to Kushtia Sadar Hospital, Kushtia, with two days of watery diarrhea and fever. The sample was collected in a container that was clean, sterile, disinfectant, and tight-fitting cover. Within two hours of collection, the collected sample was brought to the Microbiology laboratory.

### Culture-based detection

Isolation of the bacterium was performed using a culture-based methods. Firstly, the stool sample suspension was prepared by adding sterile distilled saline water. Then, the sample suspension was inoculated to Salmonella and Shigella Agar (SS Agar) media, Mac Conkey agar media, and nutrient agar media (red or pink-orange color morphology) by using a sterile cotton bud and the incubation process lasted 24 hours at 37°C. The distinct isolates (3-5 colonies) were purified by repeated sub-culturing on the same media, streaked on a nutrient agar plate, then another incubation for a full day at 37°C. The manufacturing company’s instructions were followed in the preparation of all the media utilized in the current study, and their sterility was tested. The isolated strain was stored at 4°C on agar plates for short duration but preserved in 80% (v/v) glycerol at −80°C for long-term storage for further analysis.

To identify the isolated bacteria, a range of biochemical tests such as oxidase, motility, indole, citrate, lysine decarboxylase, urease, triple sugar iron (TSI), and a range of sugars were conducted. Morphological characteristics, including colony shape and colour were also considered. MacConkey agar was used for differentiation of lactose fermenters from non-lactose fermenters. Uninoculated media were used as negative controls.

### Molecular-based detection

#### DNA extraction and 16S rRNA Gene Amplification, Sequencing, and Analysis

Genomic DNA extraction from isolated strains was performed using the Genomic DNA Extraction Kit (Promega, USA) based on the manufacturer’s guidelines. In brief, genomic DNA extraction was conducted by first mixing 400 µL of nuclease-free water, a pure bacterial colony, and 200 µL of lysis buffer in an Eppendorf tube, followed by vertexing. Then, 400 µL of the sample-containing solution was transferred to a reagent cartridge tube, and a plunger was added to the plunger hole before placing the cartridge inside the extraction machine (Promega, USA). Subsequently, 60 µL of elution buffer was added to an elution tube and also placed in the machine. After 40-45 minutes, the extraction process was completed, and the extracted DNA sample was collected. The extracted DNA’s purity and concentration were assessed using a Promega Quantus™ Fluorometer and stored at −20°C.

The amplification process was started using universal oligonucleotide primers, 27F (5’-AGAGTTTGATCCTGGCTCAG-3’) and 1492R (5’-TACGGTTACCTTGTTACGACTT-3’). A total of 25 µL reaction volume for each bacterial isolate composed of 12.5 µL OneTaq 2× Master Mix with the Standard Buffer, 1 µL for each primer, 2 µL genomic DNA, and 9.5 µL nuclease-free water were used for PCR amplification on DNA Engine DYADTM Peltier Thermal Cycler (BIO-RAD, C1000 Touch TM, Hercules, CA, USA) [24]. The PCR conditions were tuned to initial denaturation at 94°C for 5 min followed by 35 cycles of amplification. Furthermore, the denaturation temperature was adjusted to 94°C for 30s, annealing at 50°C for 30s, extension at 68°C for 1 min, and final extension at 68°C for 10 min. After amplification, PCR products were checked in 2% agarose gel prepared in 1×TAE buffer and heated in microwave for 4 min. After cooling, 10 µL ethidium bromide was added for the electrophoresis. The band size of the amplicons was measured using a 1 kb molecular marker. After that, the gel was visualized in a Chemidoc TM imaging system (BIO-RAD Laboratories, Hercules, CA, USA). DNA purification, cycle sequencing, and DNA sequencing of PCR amplicons were performed using a BDT v3 Cycle sequencing kit in a genetic analyzer 3130. PCR products were examined via agarose gel electrophoresis, purified, and sequenced using a genetic analyzer. Sequence analysis was conducted to identify the bacterial strain. All sequences were compared against entries in the NCBI BLAST database, considering parameters such as maximum score, coverage, identity, and E-value [25].

#### Antibiotic Profiling

Antibiotic susceptibility testing was conducted using 10 different antibiotic discs, including (Erythromycine (15 μg), Kanamycin (30 μg), Penicillin G (10 units), Nalidixic acid (30 μg), Polymyxin B (300 units), Ceftriaxone (30 μg), Ceftazidine (30 μg), CO-Trimoxazole (25 μg), Colistin (10 μg), and Doxycycline (30 μg). At least 3-5 well-isolated colonies from a pure culture were selected and swabbed onto Mueller-Hinton agar plates (Markey et al., 2013). Antibiotic discs were dispensed onto the inoculated plates and incubated at 37°C for 24 hours. The zone of inhibition (or zone of clearance) surrounding each antibiotic disc was measured to determine the antibiotic’s sensitivity. A ruler or caliper was used to measure the diameter of the zone of inhibition from the disc’s edge to the close millimeter. The measurements were recorded and compared with the established protocols of the CLSI [29]. Based on these parameters, each antibiotic tested was used to classify the bacterial isolates as sensitive (S), intermediate (I), or resistant (R) to each antibiotic tested.

#### Whole Genome Sequencing, Assembly, and Annotation

The genomic DNA fragments were generated from dsDNA fragments, which underwent processes including end repair, phosphorylation, and the insertion of polyA tails, followed by sequencing adapter insertion. DNA fragments were then purified using AMPure XP magnetic beads, and PCR was employed to create a sequence library with target fragments ranging from 300 to 400 bp. Subsequently, PCR products were purified again using AMPure XP magnetic beads, and their quality was assessed using an Agilent 2100 bioanalyzer. The qualified products were sequenced using the Illumina MiniSeq System platform. Following complete genome sequencing, various quality assurance tests were conducted, including FastQC analysis using Illumina Base Space sequence analysis Hub. The genome was assembled using SPAdes assembler version 3.5. The resulting assembled genome was annotated and analyzed using the NCBI Prokaryotic Genome Annotation Pipeline (PGAP) version 4.5 and Prokka.

### Phylogenetic analysis

We extracted 22 ribosomal 16S rDNA sequences and 20 whole genome sequences that were acquired from GenBank (https://www.ncbi.nlm.nih.gov/genbank/) and aligned them using MEGA 11.0 [30]. The information about genome sequences from NCBI-acquired and used in phylogenetic and comparative analysis in Supplementary Table 01. These sequences were used to construct neighbor-joining trees using MEGA [30]. This phylogenetic tree was created using the maximum likelihood and neighbor-joining trees MEGA method with a bootstrap analysis with 1000 replicates. Bar, 50 substitutions per site.

### Data deposition

The 16S rRNA sequence and assembled whole genome sequence were submitted to GenBank under accession numbers MN726679 and WMCE00000000, respectively. The information of whole genome sequence is available under the Bioproject PRJNA590029, Biosample SAMN13316584, genome sequence version WMCE00000000, respectively.

### Strain classification

Strain classification was performed through various analyses including 16S rRNA and whole-genome-based phylogenetic tree analysis, average nucleotide identity (ANI), and digital DNA-DNA hybridization (dDDH) analysis. In order to isolated strain delineation, the phylogenetic tree was created using MEGA v. 11 [31] by the maximum likelihood method based on 16S rRNA gene coding sequences and the neighbor-joining method with 1000 bootstrap replica was set up. The whole genome-based tree was constructed first on the REALPHY 1.12 online server [32] and it was modified by MEGA v. 11 software [31]. On the other hand, the JSpecies WS [33] and GGDC web server 3.0v [34] were used for ANI and dDDH analysis [35].

### Genome comparison with reference strain with other closely associated strains

The annotated whole genome sequence was compared with the five most closely related genomes to find the identity and variation picture by pangenomic analysis using GView Server[36] and D-GENIES dot plot software [37]. D-GENIES is a freestanding and web program that does massive genome alignments with the minimap2 software package and generates interactive dot plots. We used as reference strain *S. marcescens* strain KS10 (CP027798.1). The comparison of the genomes is given in **Table 1**. The used strains were *S. liquefaciens* strain FG3 (CP033893), *S. marcescens* LCT-SM213 (NZ_JH670250), *S. marcescens* strain BWH-23 (CP020501), *S. nematodiphila* strain DSM 21420 (JPUX01000001), and *S. ureilytica* strain Lr54 (NZ_JSFB01000001). Again, the study genome was compared with three more closest genomes published in NCBI using the progressive mauve software[38], and the utilized homologs were *S. marcescens* LCT-SM213 (NZ_JH670250.1), *S. nematodiphila* DSM 21420 (JPUX01000001.1), and *S. ureilytica* strain Lr54 LG59 (NZ_JSFB01000001.1). This comparison aids in identifying genetic variations and similarities among the strains.

**Table 1.** Comparison of four reference genome with *Serratia marcescens* IU-BTGE-M_3.

| **Genome features** | ***Serratia* sp.**<br>**IU-BTGE-M-3**<br>**(This study)** | ***Serratia***<br>***marcescens* strain**<br>**BWH-23** | ***Serratia marcescens***<br>***LCT-SM213*** | ***Serratia marcescens***<br>***Strain KS10*** | ***Serratia nematodiphila***<br>***DH-S01*** |
| --- | --- | --- | --- | --- | --- |
| Genome coverage | 13.0x | 564.56x | 150.0x | 400.0x | 100.0x |
| Genome size (bp) | 4,977,583 | 5,072,681 | 5,018,857 | 5,199,559 | 5,256,558 |
| GC content | 59.70% | 59.5% | 59.5% | 59.5% | 59.5% |
| Scaffold N50 | 4.8 Mb | 5.2 Mb | 4 Mb | 5.2 Mb | 5.3 Mb |
| Scaffold L50 | 1 | 1 | 1 | 1 | 1 |
| Genes (total) | 4,827 | 4,897 | 4,795 | 5,037 | 4,992 |
| Genes (coding) | 4,681 | 4,728 | 4,662 | 4,857 | 4,837 |
| CDSs (total) | 4,721 | 4,771 | 4,704 | 4,901 | 4,867 |
| CDSs (with protein) | 4,681 | 4,728 | 4,662 | 4,857 | 4,837 |
| Number of RNAs | 106 | 126 | 91 | 131 | 125 |
| rRNAs | 8 | 22 | 5 | 27 | 22 |
| tRNAs | 82 | 89 | 73 | 91 | 89 |
| Non-coding RNAs (ncRNAs) | 16 | 15 | 13 | 13 | 14 |
| Assembly | GCA_017298695.1 | GCA_900457055.1 | GCA_900457055.1 | GCA_900457055.1 | GCA_000738675.1 |
| Antibiotic resistance genes | 4 | 17 | 8 | 23 | — |

Additionally, we compared our genome (*S. marcescens* BTGE M_3) with the reference genome sequences of *S. marcescens* strain BWH-23 (GCA_900457055.1), *S. marcescens* LCT-SM213 (GCA_900457055.1), *S. marcescens* strain KS10 (GCA_900457055.1) and *S. nematodiphila* DH-S01 (GCA_000738675.1). The orthologous gene cluster analysis tool OrthoVenn3[39], was utilized to examine the predicted protein-coding sequences from the whole genome data of *S. marcescens* BTGE M_3, *S. marcescens* strain KS10, *S. marcescens* strain BWH-23, and *S. nematodiphila* DH-S01.

Gene Ontology (GO) [40] annotations revealed that *S. marcescens* IU-BTGE-M_3 -unique genes were categorized into three main GO groups-biological, molecular, and cellular processes. Gene family expansions and contractions were investigated using the CAFÉ software[41].

### Multidrug resistance gene identification

Multidrug resistance genes were identified by the ResFinder online server [42] database where the threshold value of 80% was used. This analysis helps in understanding the potential resistance profile of the isolated strains.

## Results

### Antibiotic Bioassay result

The isolated strain of *S. marcescens* IU-BTGE-M_3 was identified using both morphology- and molecular-based methods. The ribosomal 16S length of this strain was 1410 bp in length (MN726679.1). Subsequently, a bioassay was performed to evaluate the antibiotic resistance of these *Serratia* isolates. Biologically viable triplicate isolates were subjected to testing against ten different antibiotics categorized into three antibiotic groups: aminoglycoside, cephalosporin-hydrolyzing class C beta-lactamase, and tetracycline efflux MFS transporter Tet. (41). Out of the ten antibiotics tested, seven were found to exhibit resistance in the *Serratia* isolates. The antibiotic bioassay results for the *S. marcescens* isolates revealed varying levels of resistance to different antibiotics. Out of the ten tested antibiotics, seven antibiotics (70%, *n=10*) showed resistance in the *S. marcescens* isolates. We provide the specific zones of inhibition (measured in millimeters) for each antibiotic. The isolates of *S. marcescens* revealed average zones of inhibition measuring 19±0.2, 22±0.86, 13±0.45, 24±.95, 24±0.67, 11±0.08, and 14±0.03 mm to Kanamycin (30μg), Nalidixic acid (30μg), Polymyxin B (300 unites), Ceftriaxone (30μg), CO-Trimoxazole (25μg), Colistin (10μg), and Doxycycline (30μg), respectively.

### Genomic characteristics

A high-quality *S. marcescens* IU-BTGE-M_3 genome exhibited with some characteristics provided in **Table 2**. The assembled genome size was 4,977,583 bp with GC content of 59.7% with 13X coverage, based on PGAP (provided NCBI site) annotation analysis. The genome consisted of 4,827 genes among which 4,681 were coding genes. Additionally, the genome comprises of 4,721genes among which 4,681 CDSs were protein-coding sequences. A total of 106 RNAs, including tRNAs (82), rRNAs (8), and ncRNAs (16). According to RAST annotation, the arrangement of the genomes covers a total of 1483 proteins (59%) with subsystems and a total of 3215 proteins (41%) without subsystems. Moreover, 521 proteins are involved in amino acid and derivative metabolism, 648 proteins are involved in carbohydrate metabolism, 235 proteins are involved in RNA metabolism, 119 proteins are involved in DNA metabolism, 182 proteins are involved in stress response as well as 273 proteins are involved in protein metabolism which was distributed with- and without-the genome. Furthermore, 51 proteins are involved in virulence, disease, and defense. Specifically, resistance to antibiotics and toxic compounds, copper tolerance (9 genes), streptothricin antibiotic resistance (1), fosfomycin resistance (1), resistance to chromium compound (1), beta-lactamase (1) and resistance to fluoroquinolones (2), and so on (Fig. 1).

**Table 2:** Genome features and annotation statistics of *S. marcescens* IU-BTGE-M_3.

| Features | Annotation statistics |
| --- | --- |
| Genome | *Serratia marcescens* IU-BTGE-M_3 |
| Genome coverage | 13.0x |
| Genome size (bp) | 4,977,583 |
| GC content | 59.7% |
| N50 | 4.8 Mb |
| L50 | 1 |
| Genes (total) | 4,827 |
| Genes (coding) | 4,681 |
| CDSs (total) | 4,721 |
| CDSs (with protein) | 4,681 |
| Number of RNAs | 106 |
| rRNAs | 8 |
| tRNAs | 82 |
| ncRNAs | 16 |
| CRISPR Arrays | 2 |

**Fig. 1.**
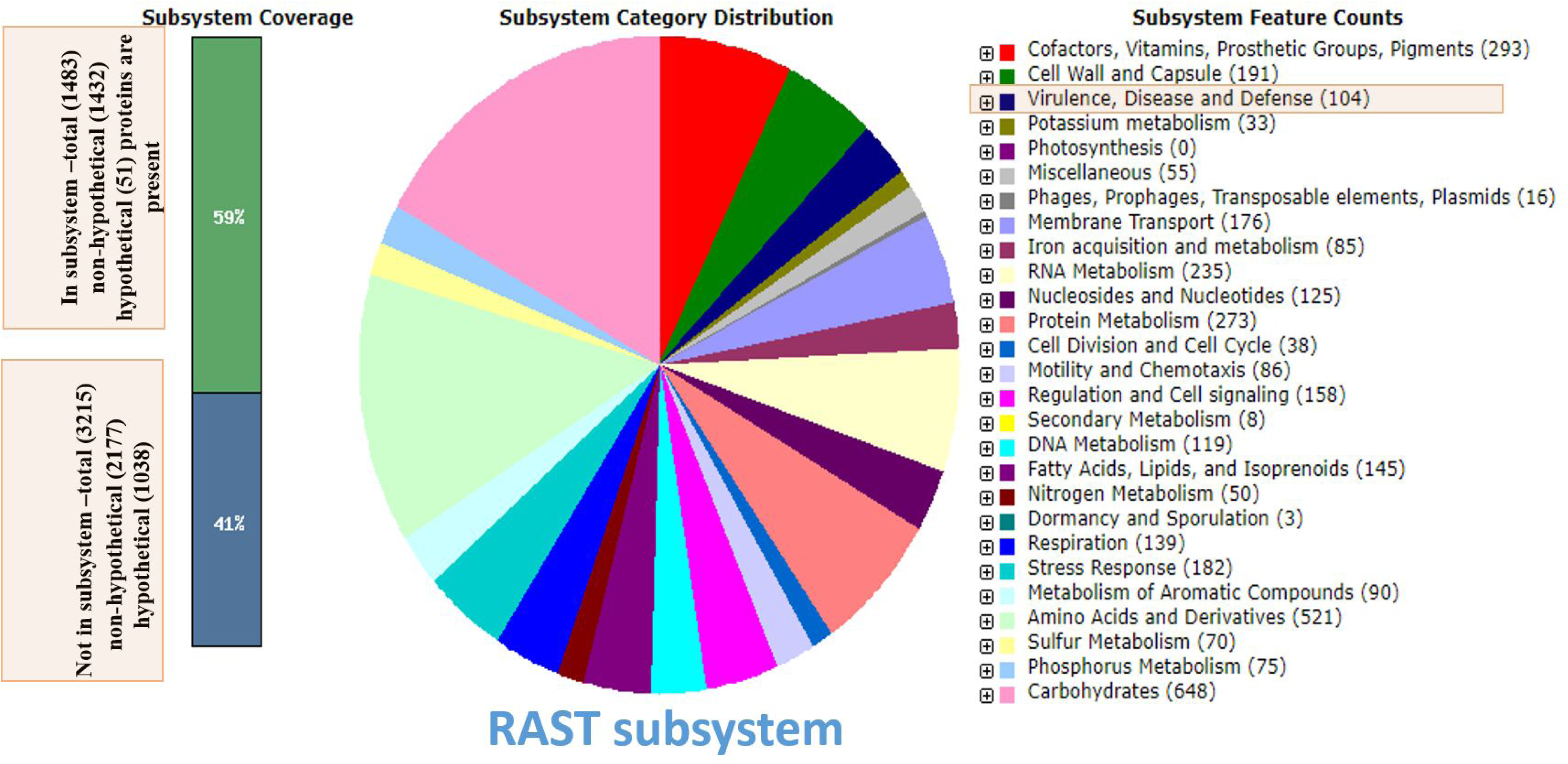
The subsystem map of *Serratia* species (*Serattia* sp. BTGE M_3) using RAST Subsystem. Genes are connected to subsystems and their distribution in different categories. The categories are expandable down to the specific gene. The virulence, disease, and defense genes are shown in blue colored. RAST server produces 70–95% of the reaction network, depending on the specific species and genome.

### Strain classification

*S. marcescens* IU*_*IU-BTGE-M_3 was closely (bootstrap value 69% similarity) related to *S. marcescens* strain KS10 while other strains were genetically distant from each other, based on the 16S-based phylogenetic analysis. Depending on the whole genome phylogenetic tree analysis, IU-BTGE-M_3 was established to be closely related to *S. marcescens* strain KS10, with >79% similarity of sequence (Fig. 2A and 2B). This conclusion was also supported by the differentiation of average nucleotide identity (ANI) and digital DNA-DNA hybridization (dDDH) results (Tables 3 and 4). The pairwise ANI blast (ANIb) values observed between the IU-BTGE-M_3 strain and *S. marcescens* strain KS10 were 98.06%, while *S. nematodiphila* DSM 21420 was 97.61%. In fact, *S. marcescens* Db11, *S. marcescens* strain BWH-23, and *S. ureilytica* strain Lr54 LG59 showed more than 90% ANIb score. Furthermore, the digital DNA-DNA hybridization (dDDH) analysis of *S. marcescens* IU-BTGE-M_3 strain with its nearest homologs *S. marcescens* strain KS10 87.5 (formula 1) and 89.9 (formula 2) and revealed GC 0.11%, whereas *S. nematodiphila* DSM 21420 showed 85% dDDH value and 0.25% GC. Where all other strains expressed 65% except *S. marcescens* subsp. sakuensis KCTC 42172 (73.5%) and *S. marcescens* ATCC 13880 (73.1%) dDDH score.

**Fig. 2.**
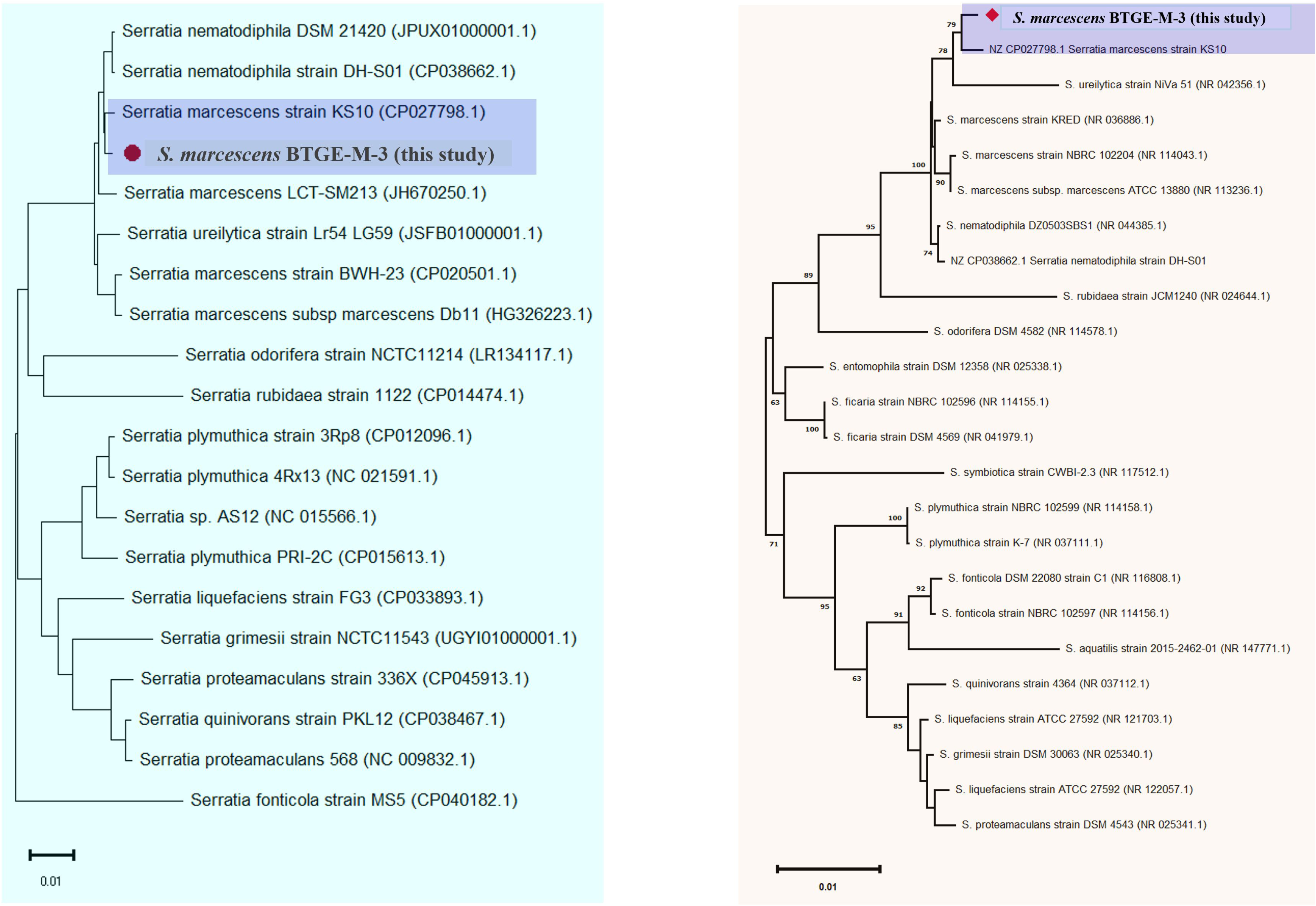
Phylogenetic Tree was constructed with the gene of 16S ribosomal (A) and whole genome sequences (B). The length of aligned gene sequence was 1580 bp. A total 24 closely similar strains were taken from NCBI. The evolutionary position of *Serattia* sp. BTGE M_3) (pointed with a red circle) was determined by using the Maximum Likelihood method based on the Tamura–Nei model on MEGA13 software. The bootstrap consensus tree was derived from 1000 replications.

**Table 3:**
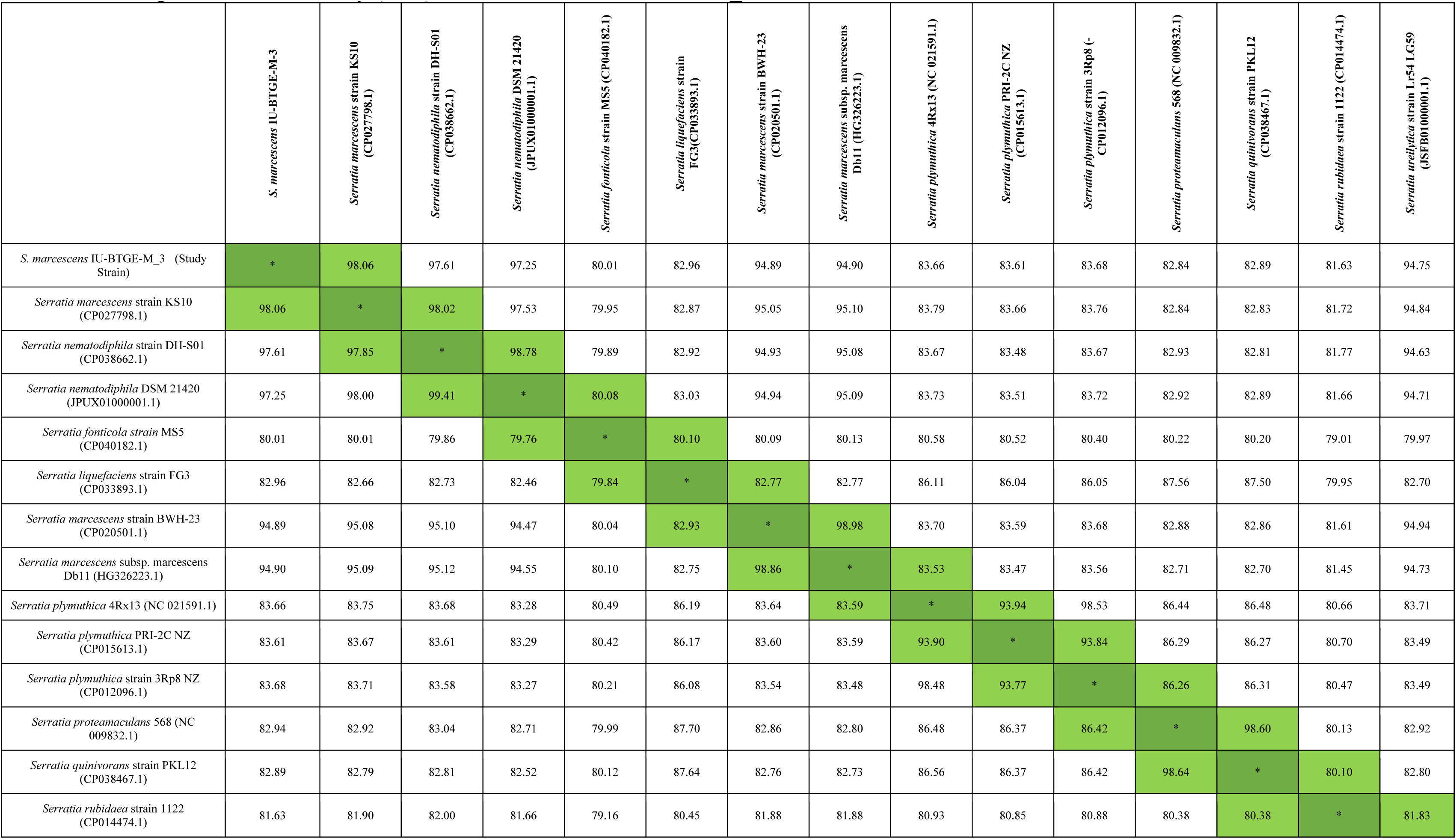
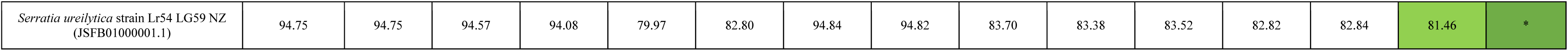
Average Nucleotide Identity (ANI) of *S. marcescens* IU-BTGE-M_3.

**Table 4:** Digital DNA-DNA Hybridization (dDDH) of *S. marcescens* IU-BTGE-M_3.

| Query genome | Reference genome strains | Formula: 1<br>(HSP length/ total length)<br>DDH | Formula: 2<br>(identities/ HSP length)<br>DDH<br>Recommended | Formula: 3<br>(identities/ total length)<br>DDH | Difference in % G+C:<br>(interpretation: distinct species) |
| --- | --- | --- | --- | --- | --- |
| *S. marcescens*<br>IU-BTGE-M_3 | *S. fonticola* strain MS5 (CP040182.1) | 29.3 | 24.5 | 27.1 | 5.84 |
|  | *S. grimesii* strain NCTC11543 (NZ_UGYI01000001.1) | 46.5 | 24.9 | 39.4 | 6.98 |
|  | *S. liquefaciens* strain FG3 (CP033893.1) | 46.8 | 27.3 | 40.7 | 4.41 |
|  | *S. marcescens* LCT-SM213 (NZ_JH670250.1) | 62.9 | 72.3 | 66.2 | 0.12 |
|  | *S. marcescens* strain BWH-23 (CP020501.1) | 84.8 | 63.3 | 83.5 | 0.32 |
|  | *S. marcescens* strain KS10 (CP027798.1) | 89.9 | **87.5** | 92.1 | 0.11 |
|  | *S. marcescens* subsp. *marcescens* Db11 (HG326223.1) | 84.1 | 63.2 | 82.9 | 0.24 |
|  | *S. nematodiphila* DSM 21420 (JPUX01000001.1) | 74.5 | 85 | 79 | 0.25 |
|  | *S. nematodiphila* strain DH-S01 (CP038662.1) | 86.7 | 85 | 89.3 | 0.22 |
|  | *S. odorifera* strain NCTC11214 (NZ_LR134117.1) | 36.7 | 25.3 | 32.9 | 3.7 |
|  | *S. plymuthica* 4Rx13 (NC_021591.1) | 50.4 | 28.4 | 43.8 | 3.5 |
|  | *S. plymuthica* PRI-2C (NZ_CP015613.1) | 46.1 | 28.6 | 40.8 | 4.09 |
|  | *S. plymuthica* strain 3Rp8 (NZ_CP012096.1) | 48 | 28.6 | 42.2 | 3.68 |
|  | *S. proteamaculans* 568 (NC_009832.1) | 48.3 | 27.2 | 41.7 | 4.68 |
|  | *S. proteamaculans* strain 336X_ (NZ_CP045913.1) | 40.4 | 27.7 | 36.3 | 4.87 |
|  | *S. quinivorans* strain PKL12 (CP038467.1) | 49 | 27.2 | 42.3 | 4.61 |
|  | *S. rubidaea* strain 1122 (NZ_CP014474.1) | 39.3 | 26.6 | 35.1 | 0.55 |
|  | *S.* sp. AS12 (NC_015566.1) | 48.1 | 28.2 | 42.1 | 3.79 |
|  | *S. ureilytica* strain Lr54 LG59 (NZ_JSFB01000001.1) | 79.3 | 62.8 | 78.8 | 0.57 |

### Comparative genome study

The local collinear blocks (LCBs) of the three genomes used for the genomic comparison did not totally match with the LCBs of the IU-BTGE-M_3 genomes. Thus, there was a genetic rearrangement of genomic regions between the two genomes in terms of collinearity. As depict in Fig. 3, the study strain shared the largest homologous region with *S. marcescens* strain KS10 (CP027798.1). The multiple genome alignment of *S. marcescens* IU-BTGE-M_3 with three *Serratia marcescens* genomes, including *Serratia marcescens* strain KS10 (CP027798.1), *S. marcescens* LCT-SM213 (NZ_JH670250.1), and *Serratia marcescens* subsp. marcescens Db11 (HG326223.1), acquired from NCBI GenBank revealed interesting insights (Supp. Table 01). In the IU-BTGE_M-3 strain genome, 30 locally collinear blocks (LCBs) were identified. These LCBs indicate regions of conserved gene order across genomes. Additionally, two major rearrangements were observed in the BTGE_M-3 genome, involving a major LCB and a segment comprising three other LCBs in inverted orientation (Fig. 3).

**Fig. 3.**
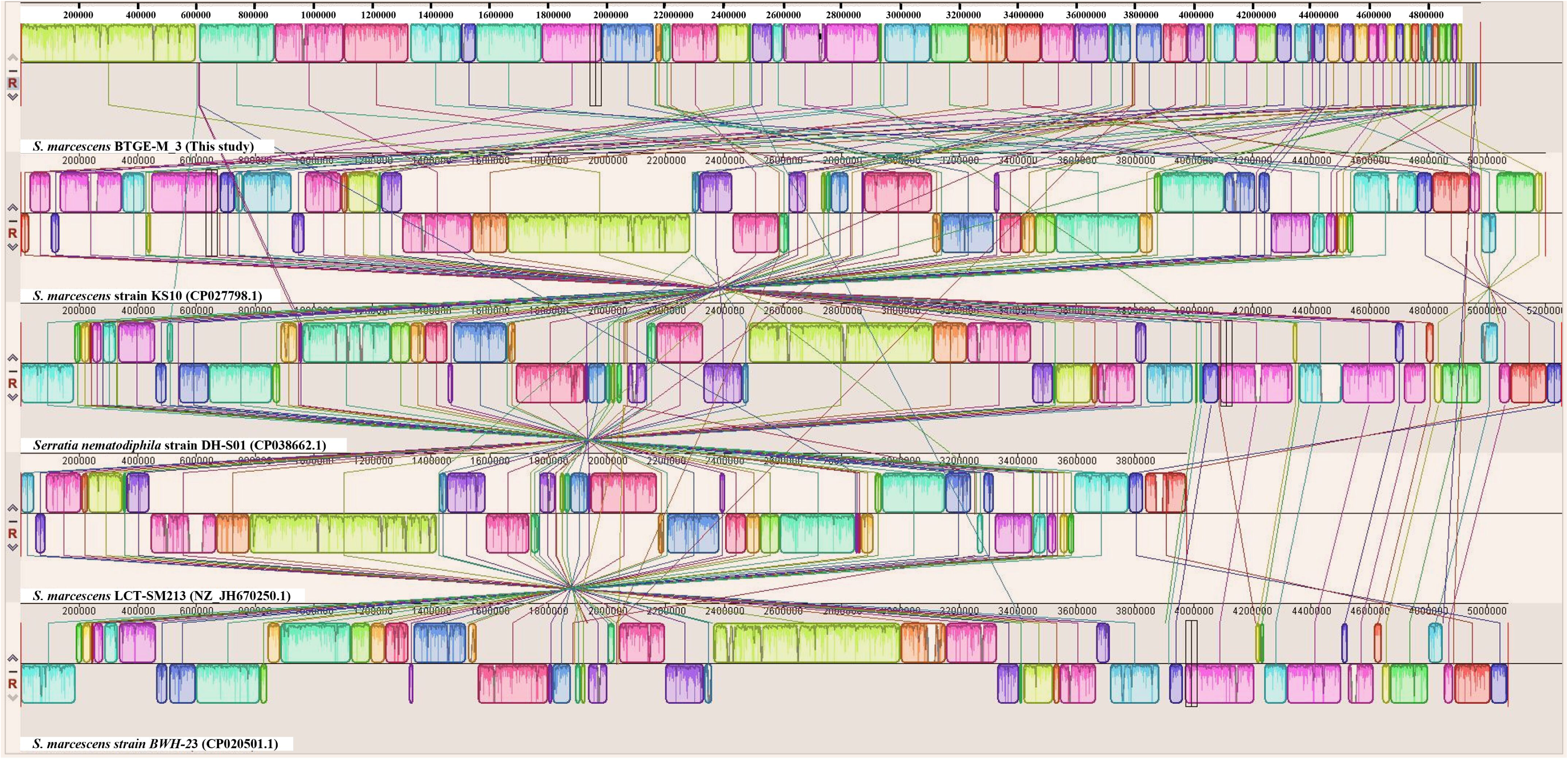
Multiple Genome Alignment with *S. marcescens* IU-BTGE-M_3. Five genomes alignment (*S. marcescens* IU-BTGE-M_3, *S. marcescens* strain KS10 (CP027798.1), *Serratia nematodiphila strain DH-S01 (CP038662.1)*, *S. marcescens* LCT-SM213 (NZ_JH670250.1), and *S. marcescens strain BWH-2*3 (CP020501.1) with MAUVE program. The figure presents the homologous regions, potential genomic rearrangements, and unique sequences within multiple aligned genomes. Each block, the software draws a similarity profile for the aligned region of the genome sequence. Completely white areas within the blocks indicate regions that were not aligned, possibly containing sequence elements specific to one genome. Colored blocks appear above and below the center line in each panel, representing regions of the genome sequence that align with parts of another genome.

Further, we performed a pan-genomic analysis of IU-BTGE-M_3 with the five closest homologs to investigate the genomic variation. The pink color slot in the circular plot represents the pangenome, while the white space denotes a region that is absent from the designated genome (Fig. 4). The circular plot clearly shows that various regions of the IU-BTGE-M_3 genome sequence were different from the other five nearest strains.

**Fig. 4.**
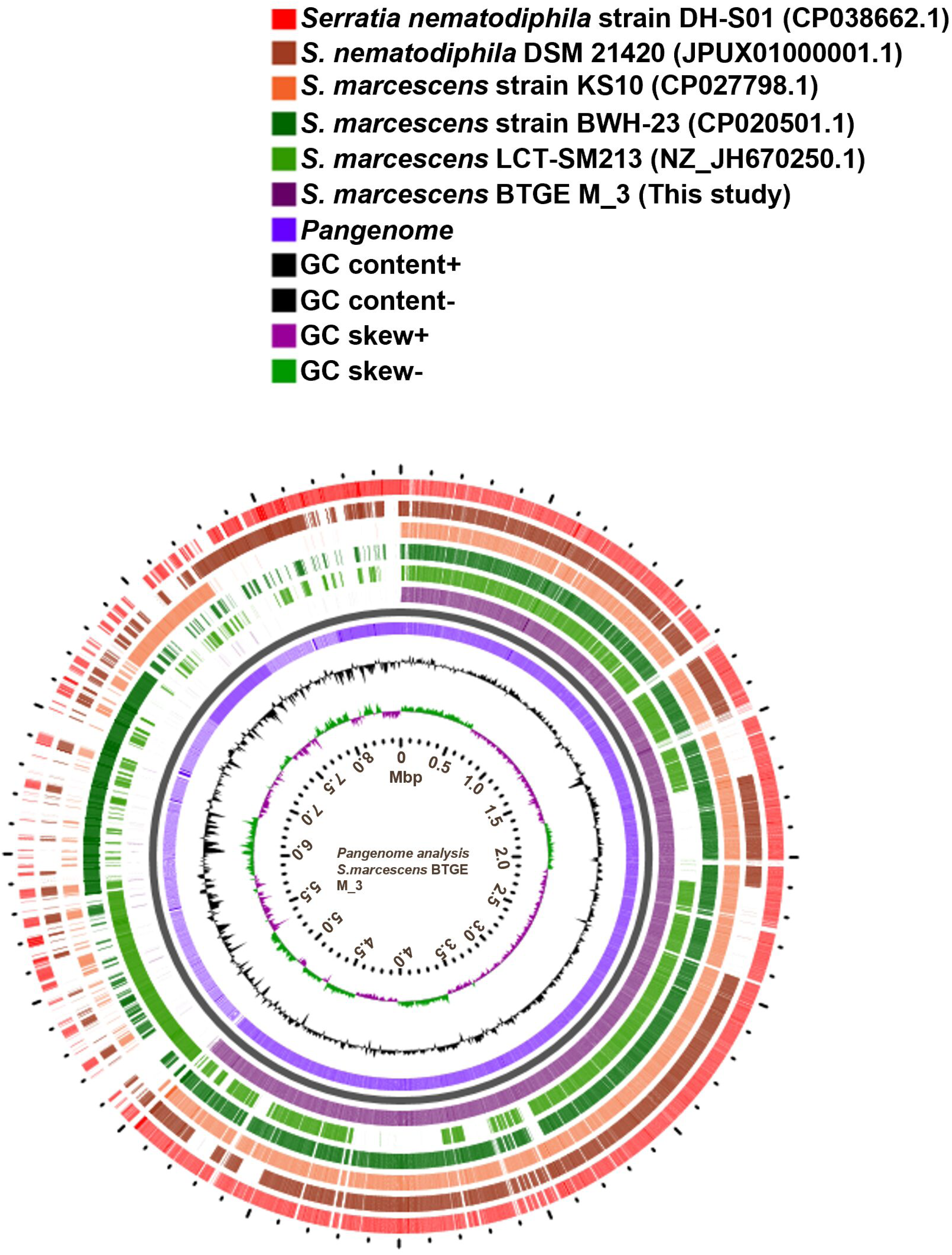
Pangenomic Analysis of *S. marcescens* BTGE M_3. Comparative study of closely related four whole genomes of *Serratia* species, including three *Serratia marcescens* (*S. marcescens* strain KS10 (CP027798.1); *S. marcescens* strain BWH-23 (CP020501.1); *S. marcescens* LCT-SM213 (NZ_JH670250.1)] and two *S. nematodiphila* strains (*Serratia nematodiphila* strain DH-S01 (CP038662.1); *S. nematodiphila* DSM 21420 (JPUX01000001.1)

Additionally, D-GENIES was used to compare the reference strain of *Serratia marcescens* strain KS10 chromosome (CP027798.1) with the strain *Serattia* sp. BTGE M_3 (Fig. 5). Visualization of the alignment of the two genomes using D-GENIES corroborates the high ANI values. Additionally, these two strains matched with comparison two genomes 86.35 (>75% similarity of sequence) while closely linked to another strain, *S. nematodiphila* DSM 21420 (72.94; >75% similarity of genome sequence) (Fig. 5). This approach provides a more realistic and comprehensive assessment of the overall similarity between the reference and query genomes that can be established only by visual inspection.

**Fig. 5.**
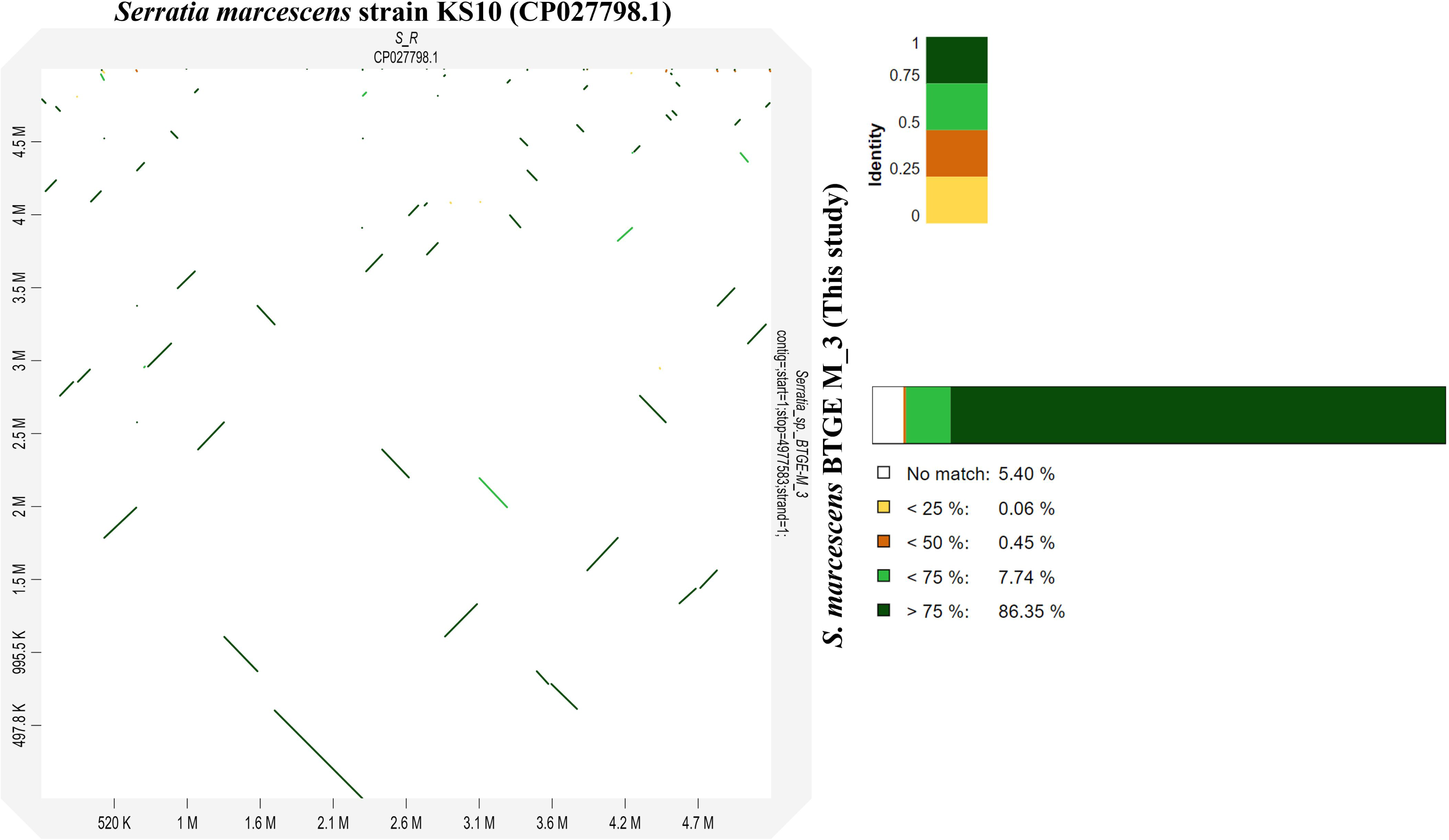
Genome comparison between the reference strain *Serratia marcescens* and *S. marcescens* BTGE M_3 using the program D-GENIES. The reference strain *S. marcescens* strain KS10 chromosome (CP027798.1) and compared strain *Serattia* sp. BTGE M_3. Sequence similarity is color coded from 0 to 1. CP027798 was used as a reference on the x-axis and BTGE M_3 as a query is on the y-axis. Genomic alignment regions are presented as five-colored lines, each corresponding to different similarity values (White: no match, Yellow: <25% similarity, Orange: 25–50% similarity, Green: 50–75% similarity, and Dark Green: >75%). The long stretches of dark green lines in the diagonal indicate high nucleotide similarity between the two strains of *S. marcescens*.

A comparative genomic analysis of *S. marcescens* IU-BTGE-M_3 with other reference genomes, specifically *S. marcescens* strain BWH-23, *S. marcescens* LCT-SM213, *S. marcescens* strain KS10, and *S. nematodiphila* DH-S01 (Fig 6). Gene Ontology (GO) annotations revealed that 271 unique gene clusters in the IU-BTGE-M_3 genome were categorized into biological, molecular, and cellular processes. Notably, some of the key biological processes included cellular metabolic processes (GO:0044464; 43.27%), antibiotic biosynthetic processes (GO:0017000; 7.39%), and molecular functions such as cyclase activity (GO:0009975) (Supporting Information, Fig. S1 and Table S2).

**Fig. 6.**
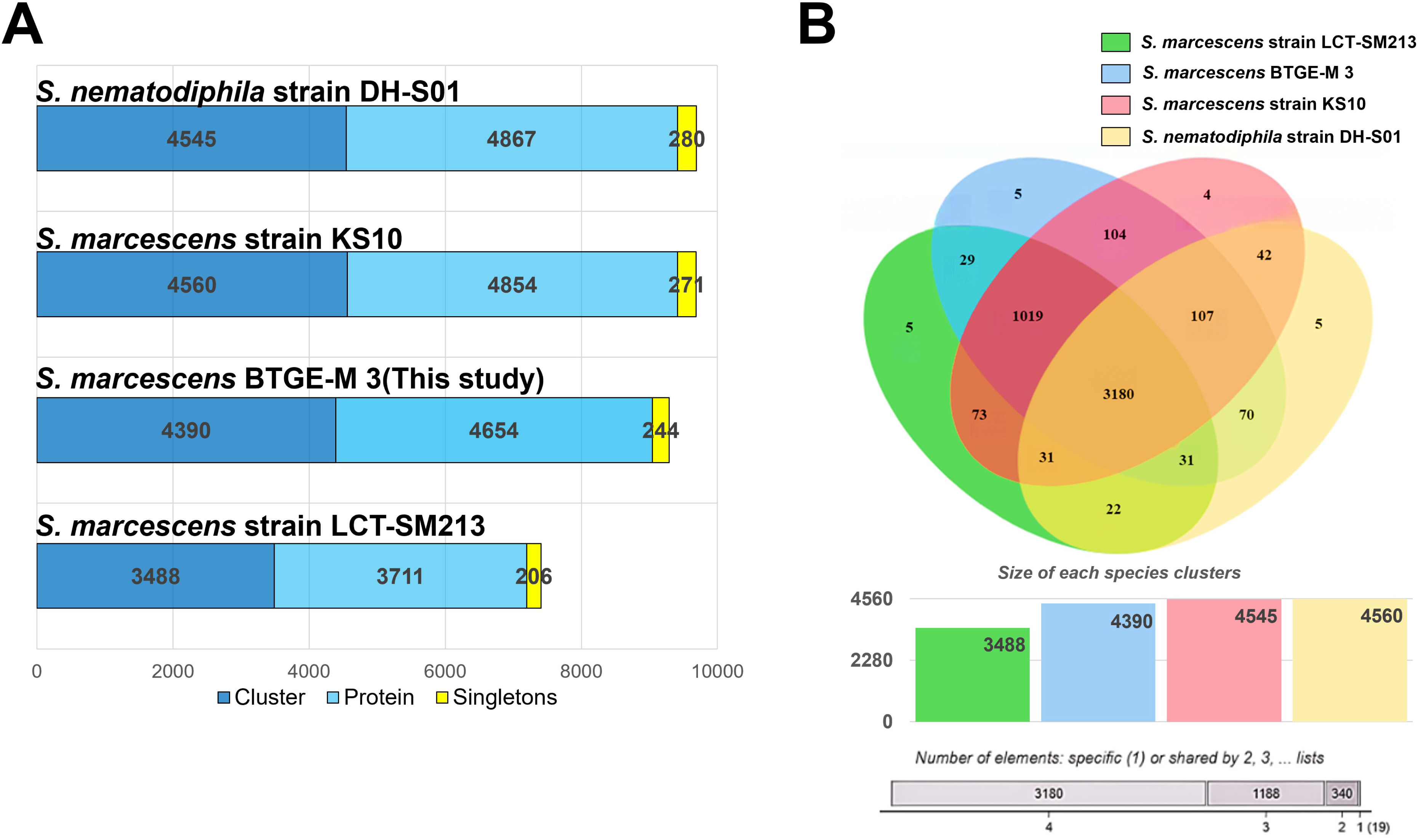
Comparison of shared gene families and orthologous clusters among the four representative *Serratia* genomes: *S. nematodiphila* strain DH-S01, *S. marcescens* strain KS10, *S. marcescens* BTGE-M 3(this study) and *S.marcescens* strain LCT-SM213. **A.** The numbers of clusters (dark blue), proteins (sky blue), and singletons (yellow) are displayed for each species in the bar chart. Singleton protein sequences refer to the sequences that did not cluster with any other sequences. **B.** The Venn diagram shows the distribution of orthologous protein clusters of predicted protein sequences, with different sections representing the number of protein clusters in each species. The graph below compares the number of clusters among the *Serratia* species.

Additionally, 32 clusters were found to be related to antibiotic resistance genes. Of particular importance were cluster 1276, which contains a tetracycline resistance protein (*tetA* gene), and cluster 3153, which includes the beta-lactamase gene (*ampC*) (Supporting Table 3).

The expansion and contraction of gene families during all evolutionary branches. The phylogenetic displayed the numbers of expanded (red) and contracted (blue) gene families at the circular shapes beside each strain name of the bacteria. Millions of years are used to express the divergence time. The IU-BTGE-M-3 strain exhibits expansions in 3 gene families and contractions in 152 gene families, respectively.

*S. marcescens* strain KS10 is much closer to other *Serratia* species. The phylogenetic reconstruction clearly supports *S. marcescens* strain KS10 other *Serratia* species (Supp Fig. 2).

### Multidrug resistance gene identification

The ResFinder web server was utilized to revealed the antimicrobial resistance genes in the isolates sequence. Genes representing resistance to four classes of antimicrobials were detected in the genomes of the isolates (Table 5). Resistance to aminoglycoside was associated with carriage of *aac (6’)-Ic* gene; resistance to fluoroquinolone was associated with carriage of *Oqx*B gene; resistance to tetracycline was associated with carriage *tet* (41) gene; and resistance to beta-lactam was associated to *blaSST-1* gene for the isolate.

**Table 5:**
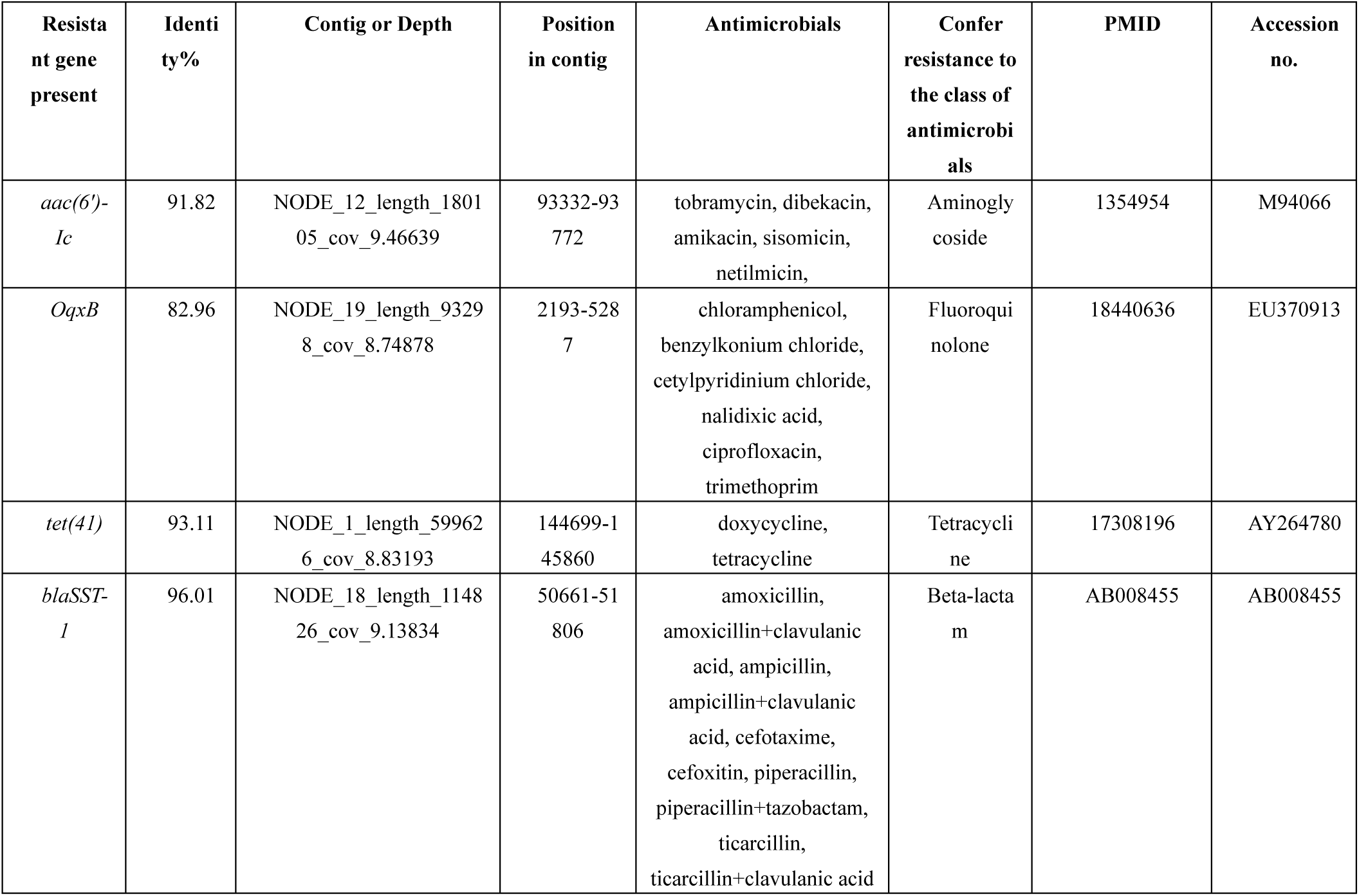
Resistance genes in the *S. marcescens* IU-BTGE-M_3 isolate detected with ResFinder.

## Discussion

The emergence of antibiotic resistance in clinical bacterial isolates, including *S. marcescens,* is a growing global issue [4,16,43]. Several prior investigations have demonstrated that *S. marcescens* strains exhibited multiple AMR genes [16], highlighting the bacterium’s resistance to a wide range of antibiotics, including beta-lactam antibiotics like penicillin, broad-spectrum antibiotics such as tetracycline, macrolides like erythromycin, aminoglycosides, fluoroquinolones, and third-generation cephalosporins, which were once commonly used to treat *S. marcescens* infections [14].

A study of 158 clinical specimens of *S. marcescens* reported resistance rates of 22.7% to ceftriaxone and 19.6% to ceftazidime. Interestingly, the resistance rates to gentamicin and cefotaxime was just 0.6% [44]. Gentamicin-resistant *S. marcescens* strains were identified in an earlier study by Graham et al. [45]. According to preliminary evidence, plasmid-mediated mechanisms are not responsible for the nonspecific aminoglycoside resistance observed [46], implying that alternative mechanisms, such as enzymatic modification, play a more significant role. A notable example is the bifunctional enzyme that has both aminoglycoside phosphotransferase (APH) and aminoglycoside acetyltransferase (AAC), [enzyme APH(2″)-AAC(6′)], which can modify nearly all aminoglycoside antibiotics, presenting a significant challenge for the development of novel aminoglycosides [47]. The current study confirmed the presence of antibiotic resistance in aminoglycoside antibiotics phenotypically and genotypically.

Recent genomic analyses have identified a multidrug-resistant *S. marcescens* isolate carrying tetracycline resistance gene, *tet* (41); aminoglycoside N-acetyltransferase gene, *aac* (6′)-*Ic*; and class C beta-lactamase-resistant gene, *bla*SST-1; all located on the chromosome. In a previous study, resistance to aminoglycosides in bacteria is typically attributed to enzymatic degradation by acetyltransferases (encoded by gene *aac*)[48]. AmpC, a chromosomal enzyme, has been discovered to deactivate carbapenem antibiotics[49] and b-lactamase, also a chromosomal gene, is expressed, leading to b-lactam hydrolysis and resistance to antibiotics such as cephalosporinase, carbapenem, and penicillinase in Enterobacteriaceae bacteria[50]. In addition, a number of virulence genes were demonstrated. The phylogenetic tree revealed that the *S. marcescens* strain from this study was comparable to other *S. marcescens* strains from different ecological niches [3]. Similarly, we revealed the four resistance genes in our isolated *S. marcescens* (Table 5). This study also detected resistance to antibiotics and toxic compounds, including copper tolerance and resistance to chromium compounds, highlighting the strain’s potential impact on clinical and environmental health.

The therapeutic management of *S. marcescens* infections is challenging due to the bacterium’s inherent resistance to multiple antibiotic classes, including ampicillin, first- and second-generation cephalosporins, macrolides, and cationic antimicrobial peptides (CAPs)[4,51]. Carbapenems or cefepime are often used to treat *S. marcescens* infections, with aminoglycoside amikacin also showing efficacy. However, resistance to tobramycin and gentamicin is on the rise [52,53]. The World Health Organization has emphasized the scarcity of treatment options for multi-resistant *S. marcescens* isolates (WHO, 2017). Recent study has also found that *S. marcescens* strains from different sources may act as repositories for antibiotic resistance determinants, focusing on the importance of ongoing surveillance and research to prevent resistance transmission[54].

In clinical settings, *S. marcescens* is often isolated from hospital environments, including nosocomial infections. For example, two infants with sepsis caused by *S. marcescens* had the strains first detected in fecal samples and later in feeding samples [55]. *S. marcescens* strain KM025385 was isolated from the pus cells of a diabetic patient [56] [9].

A study by Beye et al. (2018) suggested that a 16S rRNA sequence similarity between 95% and 98.65% with closely related species might indicate a new species. [57]. However, our results showed that the 16S rRNA gene sequence similarities between our study strain and closely related species were all above the 95% threshold, indicating a close relationship within the genus [3]. However, relying solely on the 16S rRNA tree for strain classification is challenging because 16S rRNA gene sequences are approximately 500-1500 bp long. Therefore, a whole-genome phylogenetic tree was constructed, which confirmed that the study strain was located with the same strain observed in the 16S tree [58].

The GC% of the strain DNA was used to determine the quality of the genome sequencing. A sequence’s GC% serves as an indicator of its quality, and in research, a GC% of at least 40–70% is considered acceptable [3]. In this regard, the genome of *Serratia* sp. FS14 is consist of a single circular chromosome of approximately 5.25 Mbp, with a genomic GC content of 59.46% and a total of 4,761 predicted CDSs [59]. Similarly, the strain studied in this analysis was composed of 59.7% GC content, indicating excellent sequence quality for further analysis. The genomic GC content of analyzed *Serratia* strains ranges from 50.9% to 59.6%. Among these, four *S. marcescens* strains exhibit the highest GC content, around 59%, while *S. fonticola* RB-25 shows the lowest GC content at 50.9%. Interestingly, no plasmid was identified in FS14, whereas other strains each contained one plasmid. However, no resemblance was found between the plasmids across diverse *Serratia* strains, contributing to the variation in genomic content [59].

Phylogenetic analysis of 16S rRNA genes placed *S. marcescens* IU-BTGE-M_3 in the same sister taxa as *S. marcescens* strain KS10 (CP027798.1), with a close similarity to the strains of *S. nematodiphila*.

Researchers utilize cutting-edge techniques for estimating whole-genome distances, leveraging algorithms to identify high-scoring segment pairs or maximally unique matches efficiently, forming a foundation for estimating intergenomic distances. For final confirmation to strain classification, DNA-DNA hybridization and average nucleotide identity analysis were carried out [60]. The highest dDDH score was 92.1%, and ANI value was 98.06% for *S. marcescens* strain KS10 (CP027798.1). According to the GGDC web server, a DDH value >70% indicates the strain belongs to the same species and >79% based on the formula: 2 (identities / HSP length) indicates the strain belongs to the same subspecies [61]. The researchers look at cutting-edge techniques for estimating whole-genome distances in their capacity to have similarities with DDH. High-scoring segment pairs or maximally unique matches can be rapidly identified using algorithms, which work well as a foundation for estimating intergenomic distances.

Another analysis of ANI cut-off value >96% for species declaration. As a consequence, our study results suggested that the strain IU-BTGE-M_3 belongs to *S. marcescens* strain KS10 species because both DDH and ANI values were exceeding the cut-off value and *S. marcescens* IU-BTGE-M_3 might be a new member of the *Serratia* species.

The present study also compared four nearest homologs genome sequences by progressive mauve, D-GENIES genies, and six closest homologs genome sequences by pangenome with the *S. marcescens* IU-BTGE-M_3 genome sequence. After progressive mauve, D-GENIES, and pan-genomic analysis, similar results were found and it was confirmed that the IU-BTGE-M_3 strain was also completely evolved from its nearest strains, indicating its evolutionary properties as well as *S. marcescens* IU-BTGE-M_3 might be a new member of the *Serratia* species. Abdullah-Al-Mamun *et al*. (2022) compared the four closest strains of genome LCBs with their study genome LCBs by multiple genome alignment [62]. They also compared the study strain with six different homologs by pan-genomic analysis and concluded that the study genome sequence was different from the nearest six homolog genomes. The study found the various addition, subtraction, inverted, rearrangement, or reshuffling in the genome[63].

The RAST figure depicts the annotation of a genome by connecting genes to functional roles within subsystems[64]. Once function assignments are determined, an initial resistance and virulence reconstruction is generated[65]. The goal appears to be determining which genes are part of active variants of these subsystems and then cataloguing these active variants to create a detailed estimate of the genome’s contents [64]. The proportion of genes in various genomes can be connected to functional roles within subsystems. The examples about 76% of the genes can be connected to *Escherichia coli* O157 genome[64]. Similarly, our genome the *Serattia* sp. BTGE M_3 is connected 59% to other published genome [66].

## Conclusion

The study provides preliminary insights into the multidrug-resistant (MDR) *S. marcescens* IU-BTGE-M_3 -strain using complete genome sequence analysis. Resistance gene predictions with phenotypic antimicrobial susceptibility tests, revealing the existence of multiple antibiotic resistance genes. The inherent resistance of *S. marcescens* to several antibiotic classes, including ampicillin, first-and second-generation cephalosporins, and subsequent generations, makes the therapeutic management of *S. marcescens* infections difficult. However, the exact effect of MDR mechanisms/pathway on overall drug resistance remains unclear. Furthermore, Gene Ontology (GO) annotations revealed that *S. marcescens* IU-BTGE-M_3 -gene clusters were categorized into three main GO groups-biological, molecular, and cellular processes. Comparative genomic studies revealed that *S. marcescens* IU-BTGE-M_3 -strain has a broad host range, wide geographic distribution, high genome variability, and that horizontal gene transfer likely plays a role in its evolution.

## Acknowledgments

The authors are grateful to the genome sequencing services from International Centre for Diarrheal Disease Research, Bangladesh (ICDDR,B)

---

**Supp. Table S1. The information about genome sequences from NCBI-acquired and used in phylogenetic and comparative analysis**

**Supp. Table S2**. **Information on cellular, molecular, and biological functions with their Gene Ontology (GO) number and their number of proteins.**

**Supp. Table S3. Information of specific cluster related to antibiotic resistance genes, their length, functions, and GO annotation features.**

**Supp. Figure S1**. Single copy cluster genes of *S. marcescens* IU-BTGE-M_3 classified as three major groups. A. Biological B. Molecular, and C. Cellular components. A detail information was provided in a supplementary Table S2.

**Supp. Figure S2**. Expansion and contraction gene numbers of *Serratia* species.

